# The tumor immune response is not compromised by mesenchymal stromal cells in humanized mice

**DOI:** 10.1101/699751

**Authors:** Gaël Moquin-Beaudry, Chloé Colas, Yuanyi Li, Renée Bazin, Jean V. Guimond, Elie Haddad, Christian Beauséjour

## Abstract

Therapeutic uses of mesenchymal stromal cells (MSCs) have emerged over the past decade. Yet, their effect on tumor growth remains highly debated, particularly in an immune competent environment. Here, we wanted to investigate the impact of human umbilical cord MSCs (hUC-MSCs) on tumor growth in humanized mice generated by the adoptive transfer of peripheral blood mononuclear cells (Hu-AT) or the co-transplantation of hematopoietic stem cells and human thymic tissue (Hu-BLT). Our results showed that the growth and immune rejection of engineered human tumors was not altered by the injection of hUC-MSCs in immune-deficient or humanized mice respectively. This was observed whether tumor cells were injected subcutaneously or intravenously and independently of the injection route of the hUC-MSCs. Moreover, only in Hu-BLT mice did hUC-MSCs have some effects on the tumor immune infiltrate, yet without altering tumor growth. These results demonstrate that hUC-MSCs do not promote tumor growth and neither do they prevent tumor infiltration and rejection by immune cells in humanized mice.

## INTRODUCTION

Mesenchymal stromal cells (MSCs) have emerged as interesting therapeutic tools for a number of ailments such as acute graft-versus-host-disease (GVHD) [1–3], diabetes [4], organ transplant [5], inflammatory[6], cardiovascular[7] and bone and cartilage diseases [8–10] among many others [11], which have led to over 600 completed or active clinical trials as of April 2019 according to the U.S. National Institute of Health (clinicaltrials.gov). However, despite clear guidelines provided by the International Society for Cellular Therapy for the characterization of MSCs [12], the breadth of isolation approaches and cell sources of these heterogeneous cells have led to confusing reports in the literature on the biological properties of MSCs.

Umbilical cord Warton’s Jelly has been identified as an excellent source of MSCs (hUC-MSCs) both from a logistical and a functional standpoint [13–16]. The simple, non-invasive harvesting procedure, the high proliferation potential [17–19], the high stemness [17, 20, 21] and the inability to transform into tumor-associated fibroblasts in vitro [22] make hUC-MSCs excellent candidates for off-the-shelf allogeneic cell therapies. hUC-MSCs have been demonstrated to be effective for wound healing, inflammation and disease management both in pre-clinical and clinical settings [3, 23–26]. The proposed mechanisms linked to these beneficial physiological effects implicate the ability of MSCs to home to sites of inflammation [27], deploy a strong immunomodulatory phenotype [13, 14, 28], have low intrinsic immunogenicity (low HLA-I, absence of HLA-DR, CD80 and CD86 costimulatory molecules) [29] and a capacity to promote tissue repair mostly via paracrine mechanisms [30]. Indeed, MSCs have been shown to secrete a multitude of soluble factors that contribute to their immunosuppressive capacities, such as lactate, indoleamine-pyrrole 2,3-dioxygenase, prostaglandin E2, nitrous oxide, transforming growth factor ß1, hepatocyte growth factor and interleukin 10 among others [31, 32]. These factors along with cell-to-cell contact mechanisms have been shown to inhibit the proliferation, maturation and function of B cells, T cells, NK cells, macrophages and dendritic cells and to regulate the function of regulatory T cells [33–38]. While these features can be desirable therapeutic tools, most have been associated to cancer development. Indeed, all aforementioned soluble factors have independent pro-tumoral properties [39–45] and demonstrated to contribute to the formation of an immunosuppressive microenvironment fueling cancer incidence and growth [46, 47].

However, debate is still on regarding the impact of MSCs on tumor growth. Many studies have shown pro-tumoral effects while just as many have demonstrated the opposite [48, 49]. The unavailability of good humanized animal models allowing to study the interaction between a human tumor and autologous immune cells has prevented researchers from measuring the potential impact of MSCs on tumor growth through alterations of the immune response. This should be a major concern considering the known immunomodulatory properties of MSCs. Hence, using two distinct humanized mouse models that combine genetically defined immune-naïve tumors with autologous or allogeneic immune humanization of immunodeficient mice, we here demonstrate that the injection of hUC-MSCs has negligible effect on cancer cell growth and immune rejection.

## EXPERIMENTAL PROCEDURES

### Isolation and culture of skin fibroblasts

Skin fibroblasts were isolated aseptically from either adult skin biopsy or fetal skin segments in accordance with the ethic committee from the CHU-SJ. In both cases, skin was cleaned out to preserve only the dermis and epidermis, triturated into 1-5mm^2^ pieces and digested with collagenase D (Roche) for one hour at 37°C with agitation. The whole mixture was then centrifugated at 400g for 5 minutes and washed with DMEM (Wisent) twice. The digested skin was seeded in 150 cm^2^ flasks in DMEM with 10% FBS and 0.2% primocin (Invivogen). Subsequent passages are also maintained in DMEM with 10% FBS and 0.2% primocin.

### Viral production

Lentiviral particles were produced by transfecting 2^nd^ (pPAX2) or 3^rd^ (pMDL and pRSV-Rev) generation packaging plasmids along with the vesicular stomatitis virus envelope (VSV-G) plasmid in 293T/17 cells (ATCC cat#CRL-11268) with polyethyleimine (PEI) in RPMI (Wisent) supplemented with 10% FBS. SV40ER cDNA was subcloned from pBABE SV40ER from William Hahn (Addgene #10891) into a lentiviral transfer plasmid containing a Neomycin resistance gene (SV40ER-Neo), Ha-Ras^V12^ lentiviral transfer plasmid containing a puromycin resistance gene (Ras^V12^-puro) was obtained from Francis Rodier (CHUM, Université de Montréal), hTERT lentiviral transfer plasmid was generated as previously described[50], mPlum was subcloned from pQC mPlum XI from Connie Cepko (Addgene #37355)[51] into a lentiviral transfer plasmid containing the puromycin selection gene. The firefly luciferase IRES-GFP (luc/GFP) cassette was inserted into a modified pHRSIN destination lentiviral vector. Particle-containing medium was harvested 30 hours later, filtered through 0.22µm syringe filter and concentrated by ultracentrifugation or frozen as is in −80°C freezer.

### Cellular transformation

Primary skin fibroblasts were transformed using a three successive lentiviral transduction (as described in Fig. S1). Cells were first transduced overnight with SV40ER-Neo viral particles. Three days later, 300µg/mL G418 (ThermoFisher) selection was applied and maintained until control GFP-transduced cells were eliminated. Cells were subsequently transduced with Ras^V12^-puro lentiviral particles and 2µg/mL Puromycin (ThermoFisher) selection was applied three days post-transduction. Cells were then transduced with hTERT lentiviral particles without antibiotic selection. Finally, cells were transduced overnight with mPlum viral particles. All transductions were carried in presence of 8µg/mL Polybrene (Sigma-Aldrich). Transformed cells were subsequently expanded and sorted using a FACSAriaII (BD Biosciences) in the APC channel for the expression of mPlum. For metastatic studies, cells were also transduced with luciferase-IRES-GFP viral particles and FACS sorted for mPlum^+^/GFP^+^ expression.

### Mouse immune reconstitution

All in vivo experiments were conducted in conformity with institutional committee for good laboratory practices for animal research (protocol #669). All mice used in this study were of the NOD/SCID/IL2Rγ^null^ (NSG) or NSG-SGM3 (expressing human IL3, GM-CSF and SCF) background, originally obtained from the Jackson Laboratory (Bar Harbor, ME) and housed in the animal care facility at the CHU Sainte-Justine Research Center under pathogen-free conditions in sterile ventilated racks. For adoptive transfer (Hu-AT) experiments, human adult peripheral blood was harvested from healthy donors after informed consent and immune cell isolated using Ficoll-Paque gradient (GE Healthcare). Buffy coat was harvested for PBMCs while granulocytes were isolated from the gradient pellet. Briefly, Ficoll was aspirated and the pellet was broken and resuspended in 38mL of sterile deionized water for 20 seconds for red blood cells (RBC) lysis before adding 2mL of sterile 20X PBS solution. PBMCs and granulocytes were counted and mixed at 1:1 ratio before injection into mice. 5×10^6^ PBMCs and 5×10^6^ granulocytes, for a total of 1×10^7^ WBC, were injected intraperitoneally (i.p.) in 200µL total volume after initial pilot experiments to determine the proper dose (Fig. S2). Age and sex-matched mice without i.p. injections were used as no-AT controls. In all cases, mice showing signs of GVHD were removed from analysis.

For BLT-reconstituted mice (Hu-BLT), 6-8 week old NSG mice were sublethally irradiated with 2Gy total body irradiation using a Faxitron CP-160 before surgical implantation of 1-2mm^3^ human fetal thymus under the kidney capsule and intravenous injection of CD34^+^ hematopoietic stem cells (HSC) isolated from autologous fetal liver as previously described [52]. Fetal (16 to 21 weeks) tissues were obtained after written informed consent (ethical committee of CHU Sainte-Justine, CER#2126). Hematopoietic engraftment was assessed by analyzing peripheral blood obtained from the saphenous vein. 50µL of blood was stained for flow cytometry using conjugated antibodies (mouse CD45-FITC from BD bioscience and 7-AAD, humanCD45-PE/Cy7, humanCD19-PE, humanCD3-APC and humanCD14-APC/Cy7 from Biolegend) and analyzed on LSRFortessa flow cytometer (BD Biosciences). Only mice with high reconstitution at week 8-10 (35-75% human CD45, >20% CD3) were used in this study. Age and sex-matched non-reconstituted NSG mice were used as control.

### hUC-MSC isolation, characterization and injection

Human umbilical cord-derived mesenchymal stromal cells (hUC-MSCs) used in this study were obtained from the Héma-Québec/Hospital Ste-Justine biobank (BaRCCO). hUC-MSCs were extracted from the Wharton’s jelly of two different umbilical cords (obtained from consenting mothers) using a proprietary explant culture method developed by Tissue Regeneration Therapeutics Inc. (TRT) and cultured in a chemically defined MSC culture medium (TheraPEAK MSCGM-CD, Lonza) as previously described [53, 54]. All procedures for cell expansion and banking were carried using detailed standard operating procedures developed for clinical grade MSC preparation. Each lot was made of up to 75 cryovials containing 13 x 10^6^ (lot 1, passage 1) or 7.5 x 10^6^ (lot 2, passage 2) cells. Phenotypic characterization of each lot was done by flow cytometry using the BD Stemflow^TM^ hMSC Analysis kit and the endothelial cell marker CD31 (all reagents from BD Bioscience) and showed that the 2 lots of hUC-MSC met the ISCT phenotypic criteria for MSCs (>95% positive for CD73, CD90, CD105 and <2% positive for the hematopoietic markers) and were <2% positive for CD31. The functional activity of freshly thawed hUC-MSCs from each lot was assessed in the inhibition of activated T cell proliferation assay, as also described previously [53, 54]. hUC-MSCs were thawed in RPMI (Wisent) immediately before injection in mice. Fluorescent MSCs were labelled and tracked as per manufacturer recommendations. In brief, freshly thawed cells were diluted at 1×10^6^ cells/mL and CellBrite TM NIR790 Cytoplasmic Membrane Dye (Biotium, cat.30079) was added at the concentration of 1µM and incubated at 37°C for 20 minutes with frequent agitation. Cells were then washed thrice in serum-free RPMI (Wisent) before being resuspended in cold serum-free RPMI (Wisent) for injection in mice. 1×10^6^ MSCs in 300µl were injected i.v. or 2.5×10^6^ MSCs in 200µl were injected i.p. Cell imaging was done using the Q-Lumi In Vivo imaging system (MediLumine, Montreal) with near-infrared filters (Ex.769-41nm, Em.832-37nm).

### Mouse orthotopic injections and monitoring

For subcutaneous tumor, 5×10^5^ transformed fibroblasts were injected in 100µL of RPMI (Wisent) to form a bulge under the skin of anesthetized mice previously shaved and wet with alcohol. Two injections per mouse (one on each side) were performed. In vivo growth monitoring was done twice weekly using the Q-Lumi In Vivo imaging system (MediLumine, Montreal) by fluorescent tracking of mPlum-expressing tumor cells (Ex. 562-40nm, Em. 641-75nm). Fluorescence signal was standardized internally for each picture and normalized using FIJI macros for picture processing. Analysis of tumor signal was also measured semi-manually using FIJI macros and expressed in fluorescence integrated density. Tumor growth was also characterized at sacrifice and compared to control tumors in each experiment. Normal growth is defined by tumors bigger than one standard deviation below control mean. Tumor growth inhibition (TGI) characterized as palpable, harvestable tumors smaller than one standard deviation below control mean.

Tumors were considered eliminated (TE) when unpalpable or too small to be harvested at sacrifice. For metastatic tumors, 1×10^6^ luciferase-expressing cells were slowly injected i.v. (tail vein) in 300µL of RPMI (Wisent) to avoid embolism and imaged weekly 10 minutes after the injection of 150mg/kg of D-luciferin using the Q-Lumi In Vivo imaging system without filters. Signal normalization and analysis was done automatically using FIJI macros and expressed in radiance (photons · s^-1^ · sr^-1^ · cm^-2^) integrated density (Area · mean intensity). All injections and surgical procedures were undergone under aseptic conditions in the CHU Sainte-Justine animal facility.

### Tumor histology

Tumor tissues were flash frozen on dry ice after harvest. 10µm-thick sections were made on a cryostat (Leica) and deposited on gelatinized microscopy slides and immediately fixed and permeabilized in 95% EtOH. Immunofluorescent staining was done against human CD45 (Cell signaling Rabbit #13917) and human CD8 (Biolegend, mouse 300901) with AlexaFluor 488 or 594 secondary antibodies and DAPI counterstain.

### Characterization the of tumor immune infiltrate

Tumors were excised, and blood was collected from mice at the time of sacrifice. Tumors were digested using the human Tumor Dissociation Kit and GentelMACS Octo Dissociation with heaters (Miltenyi). Cells were then filtered on 70µm MACS SmartStrainers (Miltenyi) and washed as per manufacturer’s protocol using RPMI (Wisent) with 10% FBS. Cells were then labelled with antibodies for analysis by flow cytometry of tumor infiltrating immune cells (TIILs). The following antibodies were used: humanCD3-AF700, humanCD33-BV510 and humanCD25-BV711 from Biolegend and mouseCD45-PE/Cy7, humanCD45-BUV395, humanCD19-PE/CF594, humanCD4-BB515, humanCD8-BV421, humanCD14-APC/H7, humanCD56-BV786, humanCD127-BB700, humanPD-1-BUV737 and humanTim3-PE from BD Biosciences. Blood samples were collected, red blood cells lysed using the BD Pharm Lyse lysis buffer (BD Biosciences) and cells stained with the same antibody panel. All data were acquired on BD LSRFortessa (BD Biosciences). Data analysis was done on FlowJo V10 (FlowJo, LLC) and the FlowSOM algorithm in FlowJo [55].

### Statistical analysis

Student’s t-test, one and two-way ANOVA with Šídák’s multiple comparison post-tests were done using GraphPad Prism 8.0. For subcutaneous tumors, n= number of tumors, 2 tumors per mouse. \**p*<0.05, \*\**p*<0.01, *** *p*<0.001, \*\*\*\**p*<0.0001.

## RESULTS

### The injection of hUC-MSCs does not affect the growth of subcutaneous tumors in Hu-AT mice

To evaluate if the immunomodulatory effect of hUC-MSCs has an impact on tumor growth, we first developed a humanized mouse model combining tumorigenic conversion of dermal fibroblasts derived from a healthy donor and adoptive transfer of white blood cells (WBCs) collected from the same (Auto-AT) or a different (Allo-AT) donor (Fig. 1A and S1). In brief, AT of 1×10^7^ WBC was done the day before mice received subcutaneous (s.c.) injection of 5×10^5^ tumor cells and the first of two intraperitoneal (i.p.) injection of 2.5×10^6^ hUC-MSCs (on day 0 and 17, see Fig. 1B). Their capacity to inhibit the proliferation of activated T cells in vitro is shown in Fig. S3A. Moreover, in vivo imaging confirmed that roughly one third of NIR790-labelled hUC-MSCs persisted for over 24 days following their i.p. injection in NSG-SGM3 mice (Fig. S3B). Using this model, we observed that tumor growth was delayed by both allogenic and autologous immune cells (Fig. 1C, 1D), but that hUC-MSCs did not have an impact on tumor growth (Fig. 1C-1E). Tumor size, either expressed in volume (mm^3^) or by fluorescence integrated intensity was not significantly changed at the time of sacrifice (Fig. 1F). Moreover, the proportion of mice receiving hUC-MSCs that had completely or partially eliminated tumors was comparable to that of non-injected mice (Fig. 1G). Using flow cytometry and immunofluorescence we also analyzed immune cells infiltration in partially rejected tumors available at the time of sacrifice (Fig. 1H and 1I). While we found that all tumors were massively infiltrated by immune cells, no statistical difference was found in the tumor immune infiltrate (CD45, CD3, CD8) in mice injected or not with hUC-MSCs.

**Figure 1.**
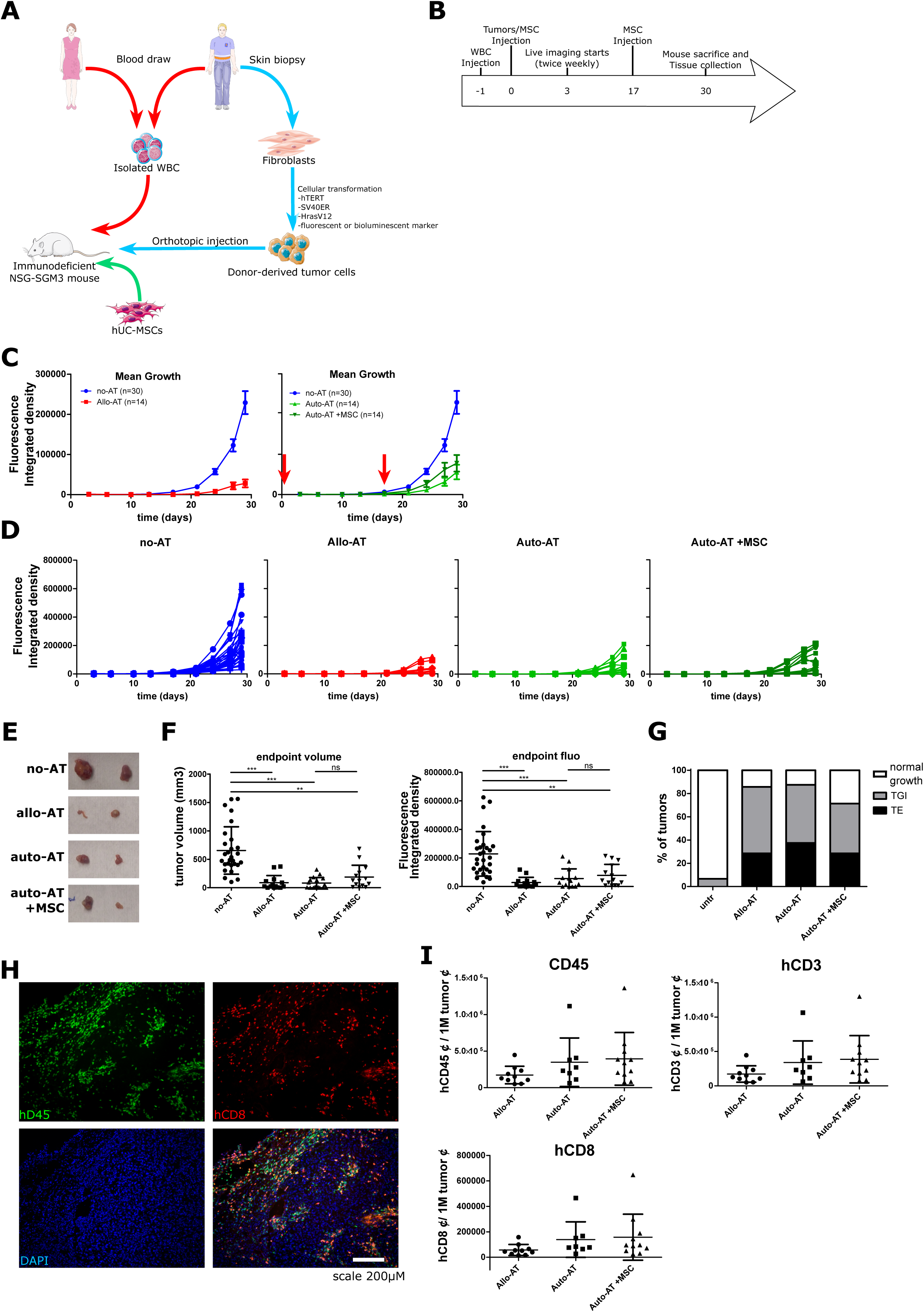
hUC-MSCs do not affect the growth of subcutaneous tumors in Hu-AT mice. (A) Schematic view of the experimental procedure and (B) Timeline of experiments. (C) Mean tumor growth without immune reconstitution (no-AT, blue) or with allogeneic adoptive transfer (Allo-AT, left panel, in red) or autologous adoptive transfer without hUC-MSCs (Auto-AT, right panel, light green) or with hUC-MSCs (Auto-AT +MSC, right panel, dark green). The red arrows indicate i.p. injection of 2.5×10^6^ MSCs. Mean±SEM. (D) Growth curve for individual tumors in the groups of panel C. (E) Representative photos of tumors for each group at sacrifice. (F) Quantification of volume (left) and fluorescence intensity (right) for all tumors at sacrifice. Each dot representing an individual tumor, mean±SD. (G) Tumor rejection assessment for each condition. TE: tumor elimination (black bar); TGI: tumor growth inhibition (grey bar) and normal tumor growth (white bar). (H) Representative immunofluorescence photos of the tumor immune infiltrate. Green: human CD45 (Top left); Red: human CD8 (Top right); Blue: DAPI (Bottom left) and merge (Bottom right). Scale 200µm. (I) Quantification of immune infiltrate for AT conditions per million tumor cells for human CD45 (Top left), CD3 (Top right) and CD8 (Bottom left). Each dot representing counts from an individual tumor.

### The injection of hUC-MSCs does not affect the growth of metastatic-like tumors in Hu-AT mice

Our observation that the i.p. injection of hUC-MSCs had no effect on the growth of s.c. tumors led us to hypothesized that a greater proximity, and perhaps cell contact, between tumors and hUC-MSCs might be necessary to impact tumor growth. To address this question, we used a metastatic-like model whereby tumor cells (1×10^6^) are first injected i.v. to colonize the lungs and liver. Within 24h, 1×10^6^ hUC-MSCs were then injected i.v. alongside the i.p. injection of 1×10^7^ autologous WBCs. Using fluorescence in vivo imaging we confirmed that more than half of the hUC-MSCs persisted over 21 days following their i.v. injection in NSG-SGM3 mice and that cells were mostly localized in the lungs and liver (Fig. S3C). Tumor burden was monitored over time by in vivo imaging and no significant variation in growth was detected between groups that received hUC-MSCs or not (Fig. 2A-C). Moreover, the injection of hUC-MSCs did not interfere with the ability of autologous immune cells to prevent/delay tumor growth in Auto-AT groups (Fig. 2A-C). When fluorescence signal intensity between lungs and liver was analyzed individually, we observed a greater reduction in tumor-associated fluorescence in lung compared to liver suggesting the AT of WBCs is more efficient at preventing or rejecting lung tumors (Fig. 2C), but this effect was also unimpaired by hUC-MSCs. Upon sacrifice of mice, these results were confirmed by the indirect quantification of tumor burden by measuring lungs and liver masses (weight and weight-to-body-weight ratio) of mice with and without auto-AT (Fig. 2D). Of note, simultaneous fluorescence imaging of tumor masses (mPlum) and labelled-hUC-MSCs confirmed that hUC-MSCs were mostly excluded from the tumor core despite their close vicinity (Fig. S3D).

**Figure 2.**
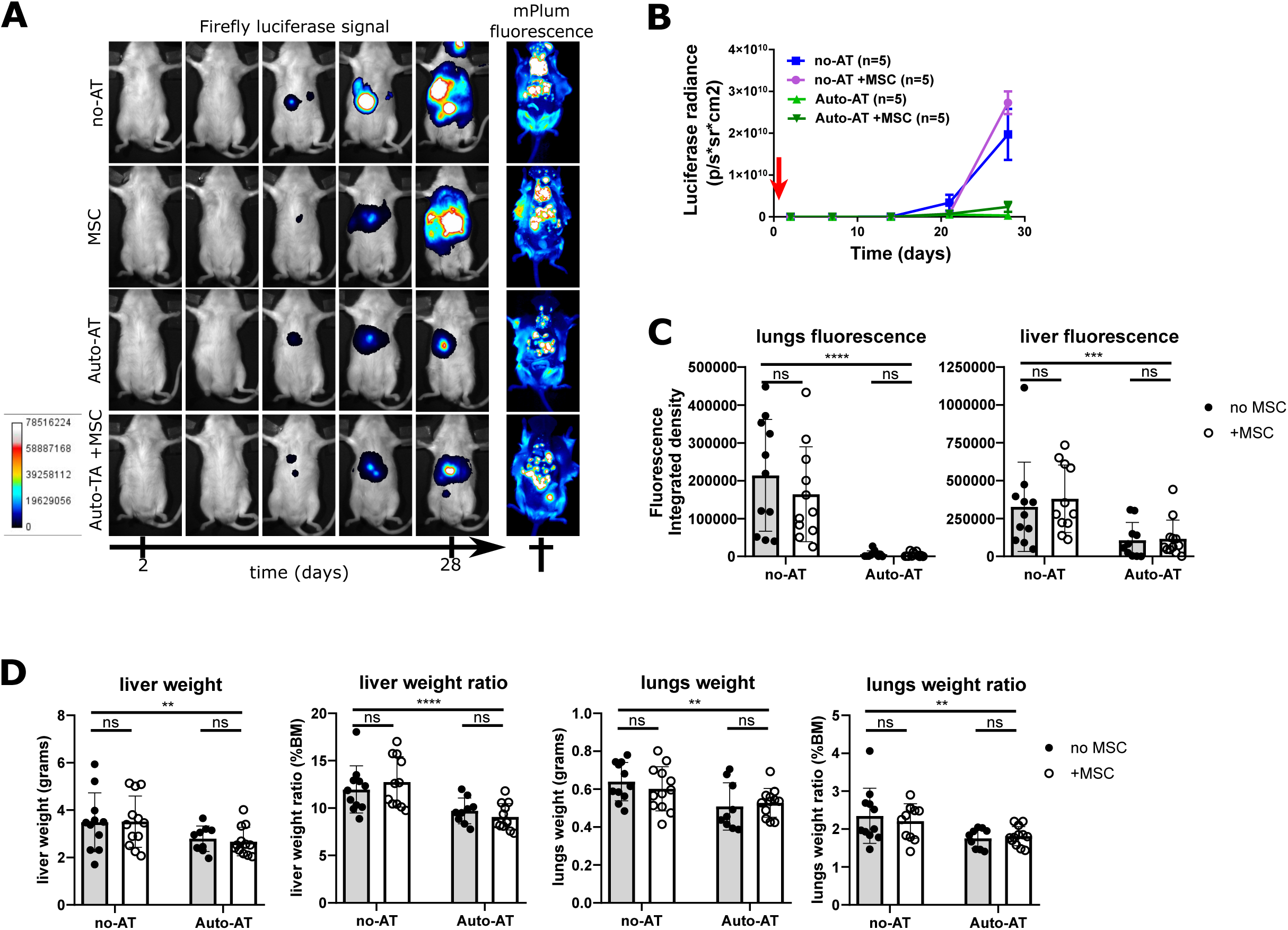
hUC-MSCs do not affect the growth of metastatic-like tumors in Hu-AT mice. (A) Tumor cells were injected i.v. and time-lapse in vivo images of lung and liver tumor growth were detected by luminescence (luciferase) in mice without immune reconstitution (no-AT), injected i.v. with 1×10^6^ MSCs (MSC), following autologous adoptive transfer of 1×10^7^ WBC (Auto-AT) or with adoptive transfer of WBC and the injection of MSCs (Auto-AT + MSC). Also shown are images of tumors by fluorescence (mPlum) at sacrifice. (B) Quantification of tumor growth luminescence over time. The red arrow indicates the i.v. injection of hUC-MSCs. n=5-6 mice per group from one of two experiments, mean±SEM. (C) Quantification of tumor size by fluorescence at sacrifice in the lungs (left) and liver (right). (D) Measurement of liver weight and weight to body mass ratio, lungs weight and weight to body mass ratio for tumors from all groups. Each dot representing the size of an individual tumor

### Human immune landscape of metastatic-like tumors is unchanged by hUC-MSCs in Hu-AT mice

To better assess whether hUC-MSCs altered the human immune landscape in mice, a 13-color flow cytometry panel was designed to characterize the circulating and tumor-infiltrating human immune populations. Manual gating of t-distributed stochastic embedding (tSNE) dimensionality reduction plots of total human CD45^+^ cells allowed the identification of two major T cell populations as shown by most cells expressing CD3 and CD4 or CD8 (Fig. 3A and C). The FlowSOM unsupervised clustering algorithm allowed for the automatic identification of 10 metaclusters based on expression of cell surface marker (Fig 3A and S4B for detailed expression). While a clear distinction between blood and tumor-infiltrating immune cells (TIIC) cluster distribution can be observed, we found no distinction between and within metaclusters in hUC-MSCs-treated animals (Fig. 3B and S4C). Further analysis revealed an enrichment in tumor infiltrating compared to circulating CD3^+^CD4^+^PD-1^hi^Tim3^hi^ dysfunctional CD4 T cells (FlowSOM.pop2 red cluster, Fig. 3B and 3C), CD3^+^CD8^+^hPD-1^hi^Tim3^hi^ dysfunctional CD8 T cells (outer sections of FlowSOM.pop0 blue cluster, Fig. 3B and 3C) and CD3^+^CD4^+^hCD25^+^CD127-regulatory T cells (outer rim of FlowSOM.pop2 red cluster, Fig. 3B and 3C). Of note, is the near complete absence of CD56^+^ clusters (FlowSOM.pop5, 8 dark green and teal clusters, Fig. 3B and 3C) in TIIC. Quantification also showed that hUC-MSCs induced no significant increase in the proportion of circulating CD4 or CD8 T cell subpopulations in blood or TIICs (Fig. 3D). However, we observed a marked increase in the proportion of T regulatory cells in TIICs compared to blood regardless of the injection of hUC-MSCs (Fig. 3D). The proportion of dysfunctional T cells (CD3^+^hPD-1^+^Tim3^+^) was also unchanged following the injection of hUC-MSCs in both blood and TIICs with no bias towards either CD4 or CD8 (Fig. 3D and data not shown). However, PD-1 and Tim3 expression levels on CD3^+^ cells were higher in tumors than in blood, with or without the presence of hUC-MSCs (Fig. 3D). Overall these results demonstrate that the immune cells landscape is greatly altered by the tumor environment but not by the injections of hUC-MSCs.

**Figure 3.**
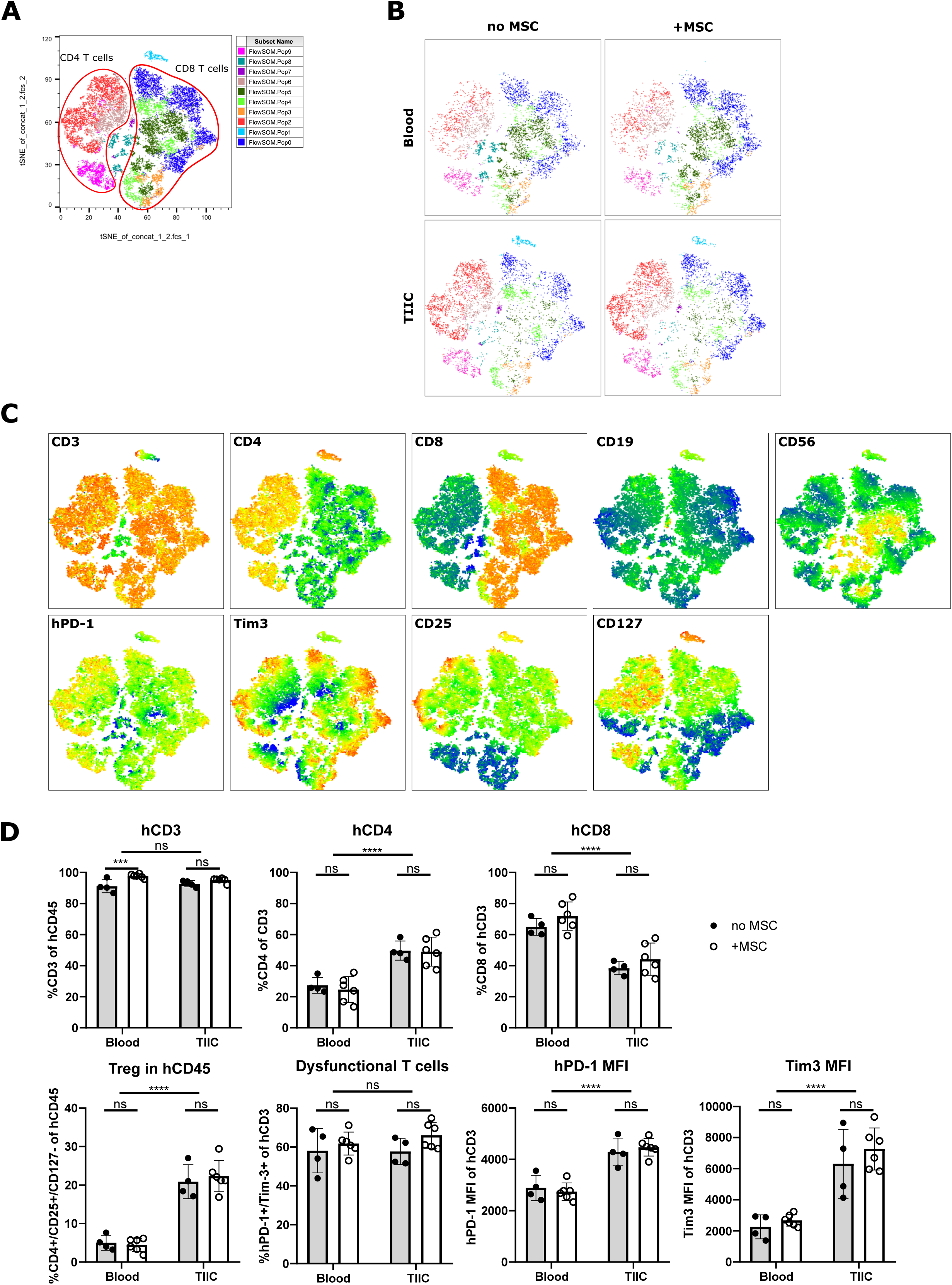
Human immune landscape of metastatic-like tumors is unchanged by the injection of hUC-MSCs in autologous Hu-AT mice. (A) Annotated tSNE plot of total human CD45 cells with unsupervised population clustering (color map) of flow cytometry data. (B) Population distribution by origin (Blood vs TIIC) and treatment (no MSC vs MSC) using the same color mapping as in (A). (C) Expression heatmap for cellular populations (CD3, CD4, CD8, CD19, CD56) (top) and phenotype (hPD-1, Tim3, CD25, CD127) (bottom) from hCD45^+^ cells. (D) Quantification of CD3 frequency in hCD45, CD4 in CD3, CD8 in CD3, CD4^+^CD25^+^CD127^-^ T reg cells in hCD45, PD-1^+^Tim3^+^ exhausted/dysfunctional cells in CD3, PD-1 and Tim3 expression level by mean fluorescence intensity (MFI). Full circles/grey columns: without MSCs; hollow circles/white columns: with MSCs, mean±SD. n=1 tumor per mouse with 4-6 mice per group were analyzed.

### The injection of hUC-MSCs does not affect the growth of tumors in Hu-BLT mice

To confirm the results obtained in Hu-AT mice, we next setup a distinct humanized tumor-immune model by using Hu-BLT mice which allows for a robust, diversified and continuous renewing of functionally mature T cells compared to what is observed following AT [56, 57]. Briefly, Hu-BLT mice were generated by the co-transplantation of human fetal liver hematopoietic stem cells along with autologous fetal thymus tissue in NSG mice (Fig.4A). Using this model, we prepared tumorigenic cell lines from fetal skin fibroblasts derived from two healthy donors (herein referred as tumor A and B, see Fig.4A). Of note, despite using the same set of defined oncogenes to generate these two cell lines, resulting tumors had different in vivo phenotypes suggesting cells were further transformed during in vitro expansion. We first observed that following the s.c. injection of 5×10^5^ cells, tumor A had a slow growth rate, as determined by in vivo imaging using fluorescence integrated density, and was eliminated in Hu-BLT mice (Fig. 4B). On the other hand, tumor B showed a faster growth rate and was not fully rejected despite being massively infiltrated by human immune cells (Fig. 4C and Fig. 5). Independently of the donor, the i.p. injection of 2.5 million hUC-MSCs (on day 0 and 14 after tumor inoculation) had no significant effect on tumor growth and the immune-rejection rate (Fig 4B and 4C). These results were confirmed by the direct quantification of tumor masses at the time of sacrifice (Fig. 4D). Similarly, the proportion of mice that had completely or partially eliminated tumors in Hu-BLT mice was unchanged by the injection of hUC-MSCs and this for both tumors (Fig. 4E). Overall, our results in Hu-BLT mice confirm our observations in Hu-AT mice and again suggest the injection of hUC-MSCs does not promote tumor growth nor does it interfere with their immune rejection.

**Figure 4.**
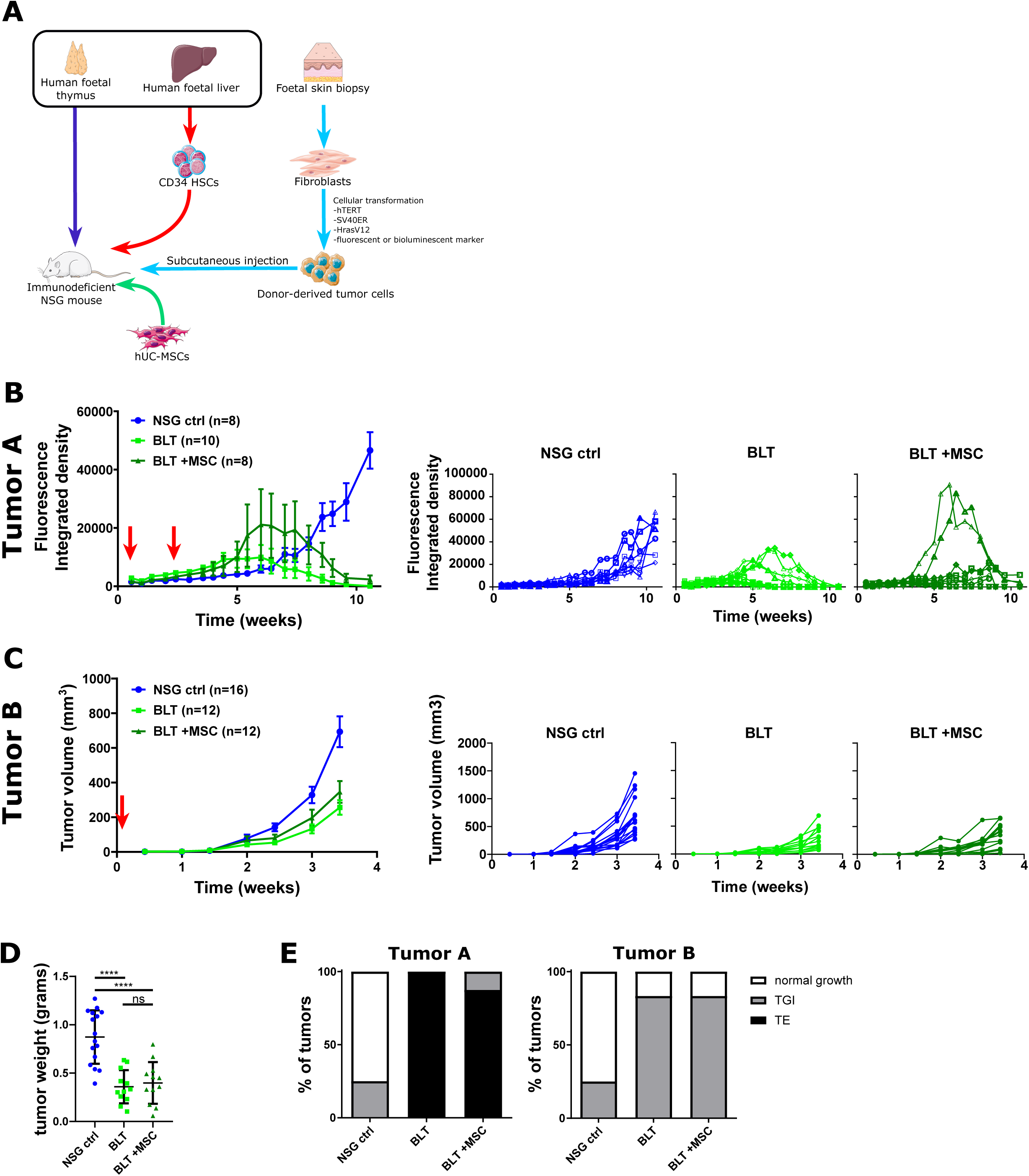
hUC-MSCs impact the immune infiltrate but not tumor growth in hu-BLT mice. (A) Schematic of experimental design. (B) Mean±SEM (left panel) and individual (right panels) tumor growth curves for tumor A without immune reconstitution (NSG ctrl), with BLT reconstitution (BLT) or with BLT reconstitution and the i.p. injection of 2.5×10^6^ hUC-MSCs (BLT +MSC). Red arrows indicate hUC-MSCs injections. (C) Same as in (B) but using a second tumor cell line (tumor B) and BLT donor. Of note, because tumor B grew faster, hUC-MSCs were injected only once. (D) Quantification of tumor weight at sacrifice. Each dot representing an individual tumor, mean±SD. (E) Tumor rejection assessment for each condition. TE: tumor elimination (black bar); TGI: tumor growth inhibition (grey bar) and normal tumor growth (white bar).

**Figure 5.**
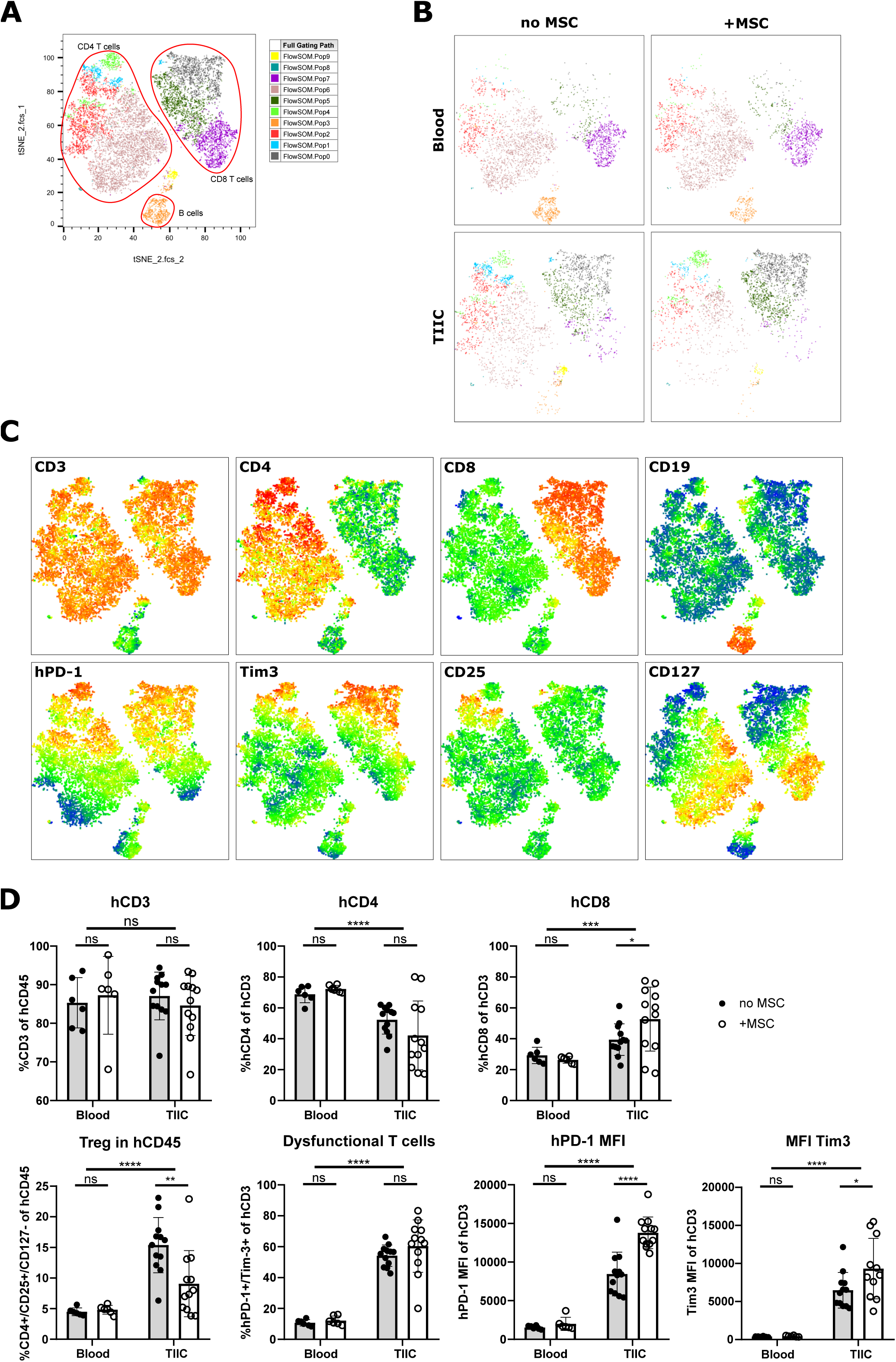
Effect of hUC-MSCs on the human immune landscape of tumors in Hu-BLT mice. (A) Annotated tSNE plot of total human CD45 cells with unsupervised population clustering (color map) of flow cytometry data. (B) Population distribution by origin (Blood vs TIIC) and treatment (no MSC vs MSC) same color mapping as in (A). (C) Expression heatmap for cellular populations (CD3, CD4, CD8, CD19) (top) and phenotype (hPD-1, Tim3, CD25, CD127) (bottom) of hCD45^+^ cells. (D) Quantification of CD3 frequency in hCD45, CD4 in CD3, CD8 in CD3, CD4^+^CD25^+^CD127^-^ Treg cells in hCD45, PD-1^+^Tim3^+^ exhausted/dysfunctional cells in CD3 and PD-1 and Tim3 expression levels by mean fluorescence intensity (MFI). Full circles/grey columns: without MSCs; hollow circles/white columns: with MSCs. Mean±SD.

### Human immune landscape of tumors in Hu-BLT mice

To assess whether hUC-MSCs altered the human immune landscape in Hu-BLT mice, we used our 13-color cytometry panel combined with tSNE analysis to identify critical populations variations in blood or TIICs. Hu-BLT mice allowed for the identification of a third defined cell cluster (B cells) in addition to the CD4 and CD8 T cells clusters, which were the only ones present in Hu-AT models (see Fig. 3A and 5A). At time of sacrifice (15 weeks post immune reconstitution), no CD33^+^, CD14^+^ or CD56^+^ populations were detected in Hu-BLT mice (Fig. S5). Cell distribution between blood and TIICs was different as shown by the near absence of B cells in tumor samples and variations in the distribution of T cell populations (Fig. 5B). For example, we observed an enrichment in the dysfunction markers PD-1 and Tim3, an increase of CD25 and a reduction of CD127 in TIICs (Fig.5B and C). Moreover, while the distribution of immune cells was mostly unaffected by the injection of hUC-MSCs in blood, we observed some significant variations in TIICs (Fig. 5B, 5D and S6B). For instance, a significant increase in the proportion of CD8^+^ T cells, but not CD4^+^ T cells, was observed within TIICs in the presence of hUC-MSCs (Fig. 5D). Similarly, regulatory T cells (CD4^+^CD25^+^CD127^-^) were significantly decreased within TIICs in the presence of hUC-MSCs (Fig. 5D). Finally, while the proportion of dysfunctional T cells (CD3^+^PD-1^+^Tim3^+^) was higher in TIICs than in peripheral blood, the injection of hUC-MSCs had no impact on this population (Fig. 5D). However, we observed that mice injected with hUC-MSC exhibited a significant increase in PD-1 and Tim3 expression in the hCD3^+^ compartment, a phenotype that was mostly observed in CD4 T cells (Fig. 5D and data not shown). Overall these results demonstrate that the immune cells landscape is somewhat altered by the injections of hUC-MSCs in Hu-BLT mice.

## DISCUSSION

The effect of MSCs on tumor growth has not reached a consensus with many studies showing contradictory results [48]. In fact, in most cases where MSCs were shown to promote human tumor growth, they were co-injected s.c. with tumor cells in an immune-deficient host and thus stimulated growth independently of their immunomodulatory properties. Instead, in a clinical setting, MSCs are likely to be injected in immune competent hosts either locally or systemically. For this reason, we thought it would be more relevant to evaluate the impact MSCs may have on tumor growth when injected at a distinct site from tumors cells and in immune competent mice. To this end, we developed two humanized mouse models (Hu-AT and Hu-BLT) using engineered naïve tumors that we showed are infiltrated and either totally or partially rejected by autologous or allogenic immune cells. We speculate some tumors were not fully rejected because of their growth kinetic, with the fast-growing tumors being more resistant (Fig 4B and C). While this remains to be demonstrated, differences in tumor growth rate may indicates that secondary mutations have occurred during the in vitro expansion period, leading to the emergence of more resistant tumor cell subpopulations. Of note, in our Hu-AT mouse model we observed better immunological clearance of lung compared to liver metastatic tumors. The reason for this is unknown but could be the result of more immune cells reaching the lungs immediately after their injection.

One advantage of our humanized autologous tumor-immune cancer models is that it can be derived easily from any healthy donor cells or tissues. Moreover, we believe our models to be more appropriate than patient-derived xenograft (PDX) to study the immunomodulatory properties of MSCs as PDX tumors were shown to be already resistant to autologous or partially matched immune cells [58, 59]. Our model thus allows to study changes in tumor immunogenicity or immune functions in an autologous setting and study mechanisms of immune-editing. Although we have not investigated the landscape of the neoantigens recognized by autologous T cells in our models, it is likely that exogenous neoantigens such as SV40 large and small T, mPlum or luciferase proteins could have fueled an HLA-I-mediated immune response [60]. Alternatively, the transformation process may have activated the expression of noncoding regions acting as tumor-specific antigens as recently described [61].

In this study we observed that the injection hUC-MSCs (either i.p. or i.v.) had no effect on tumor growth in both of our humanized models. This was surprising given the robust in vitro immunosuppressive capabilities of the hUC-MSCs used in this study (Fig. S3A). We believe this effect is due to the already suppressive tumor microenvironment in emerging tumors to which the contribution of hUC-MSCs is redundant. This absence of effect was also observed with a different hUC-MSC donor (data not shown), suggesting this effect is not donor-dependent. Doses of injected hUC-MSCs is also unlikely to be responsible for these results as the doses used in this study were considerably higher than ones commonly used clinically (16-83 times higher). Indeed, while patient i.v. dosage typically ranges from 1-2×10^6^ MSCs/kg [1, 62], we used an equivalent of 3 or 8×10^7^ hUC-MSCs/kg depending if cells were injected i.p. or i.v. respectively.

In depth analysis of tumor infiltrating cells using flow cytometry and tSNE dimensional reduction plots revealed a marked difference between circulating and TIIC profiles with for example a higher proportion of Tregs and dysfunctional T cells in tumors vs blood both in Hu-AT and Hu-BLT models (Fig. 3D and Fig.5D). Surprisingly, the injection of hUC-MSCs had almost no effect on the tumor immune infiltrate with cluster distribution and population overlap being very similar between groups. However, only in the Hu-BLT model did the injection of hUC-MSC result in an increase in infiltrating CD8 T cell frequency combined with a decrease in the proportion of Tregs (Fig 5D). Such a decrease in Tregs was very surprising given MSCs were previously shown to increase Treg recruitment and function [15, 63–66]. However, this decrease was still insufficient to alter the tumor growth profile of hUC-MSC recipients. Similarly, while the injection of hUC-MSCs had no effect on the proportion of dysfunctional T cells, we observed an increase in the expression levels of PD-1 and Tim3 within T cells in Hu-BLT mice (Fig. 5D), although this was attributable to variations in the CD4+ and not CD8+ compartment (data not shown), suggesting that CD8 T cytotoxic effector cells were globally unaffected by hUC-MSCs. While both mouse models have shown convincing tumor rejection capabilities, differences in the immune cells source and reconstitution are likely responsible for the above described dissimilarities.

One limitation of our mouse models is that not all human immune cells were adequately represented with for examples macrophages, dendritic, NK and myeloid cells being virtually absent. Reconstitution in humanized mice also does not support robust myeloid and NK cell engraftment in absence of human cytokines such as IL-15[67]. Despite the injection of both PBMCs and granulocytes in Hu-AT mice, persistence of immune cell types other than T cells was very low or undetectable at time of sacrifice. While some circulating CD14+ monocytes were generally present in Hu-BLT mice, these cells were not detected within tumors.

Overall, our study demonstrated that the injection of hUC-MSC has a minor impact on the tumor immune landscape with no significant consequences on tumor growth in humanized mouse models. These results suggest that transient therapies using hUC-MSCs are unlikely to increase cancer risk in patients.

## ACKNOWLEDGMENTS

We are grateful to the flow cytometry and animal facility for providing technical support and to Renée Dicaire for handling clinical samples. This work was supported by a grant from la Fondation Charles Bruneau to C.M.B. and E. H. G.M.B. was supported by a studentship from the Canadian Institute of Health Research.

## CONFILCT OF INTEREST STATEMENT

The authors declare no competing interests.

