## Supplementary material for "The tumor immune response is not compromised by mesenchymal stromal cells in humanized mice": supp figures

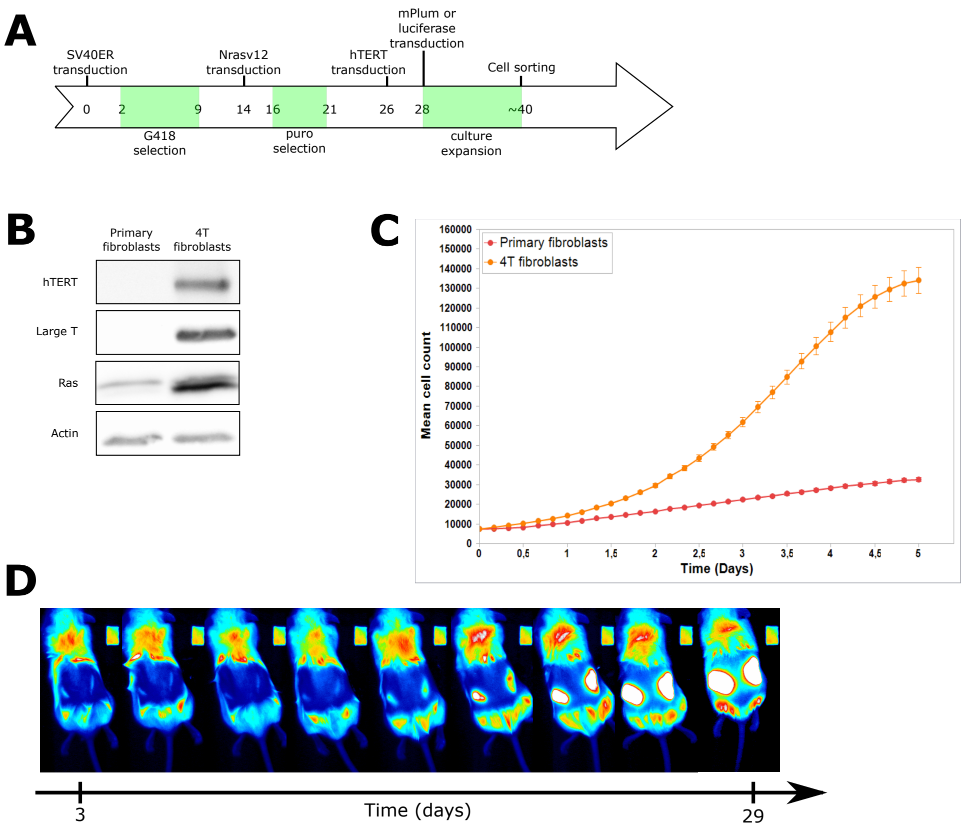


**Fig S1. Primary fibroblast transformation process.** (A) Lentivirus-based transformation protocol timeline. (B) Validation of transgene expression by Western Blot. 4T fibroblasts: fibroblast population expressing all 4 transgenes. (C) Differential *in vitro* growth curves of primary and transformed 4T fibroblasts. (D) Sample time-lapse *in vivo* imaging of tumor formation in NSG mice

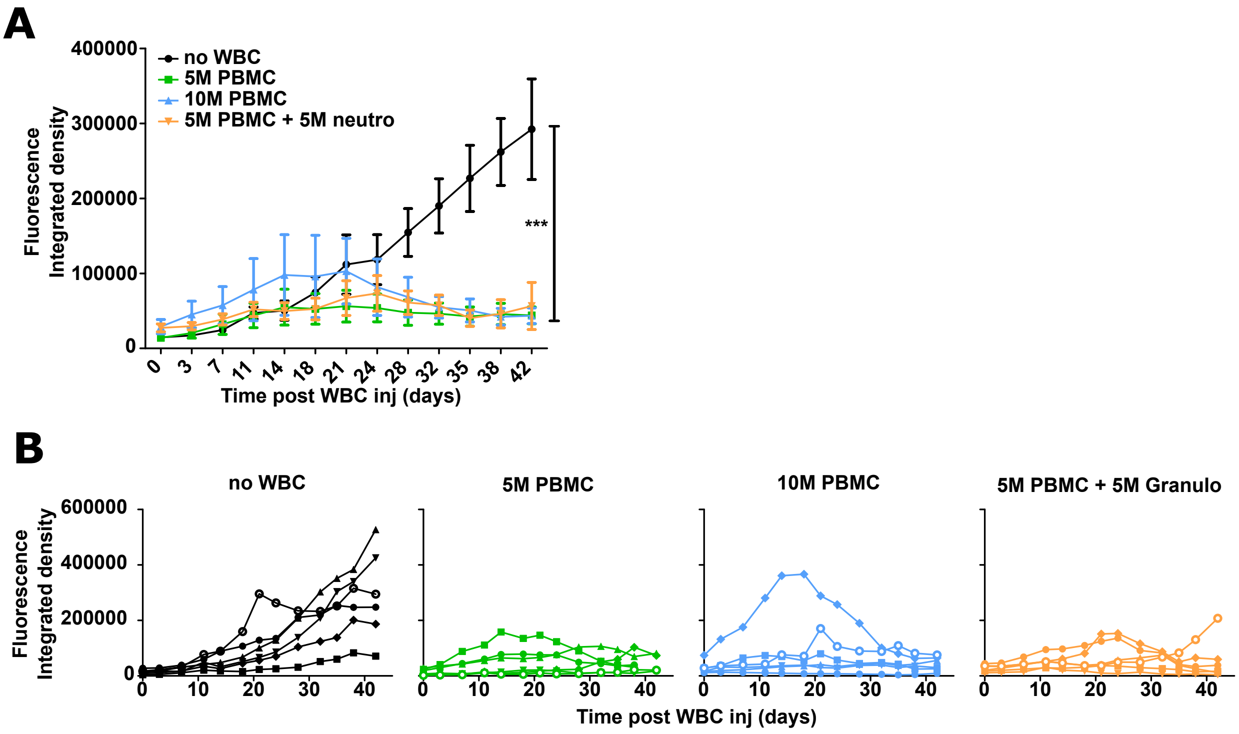


**Fig S2. WBC dosage for AT experiments.** (A) Mean±SEM and (B) detailed growth curves for different WBC regimens to assess their tumor rejection efficacy. Black: without AT; green: $5x10^6$ PBMCs; light blue $10^7$ PBMCs; orange: $5x10^6$ PBMCs + $5x10^6$ granulocytes. n=6 tumors/group.

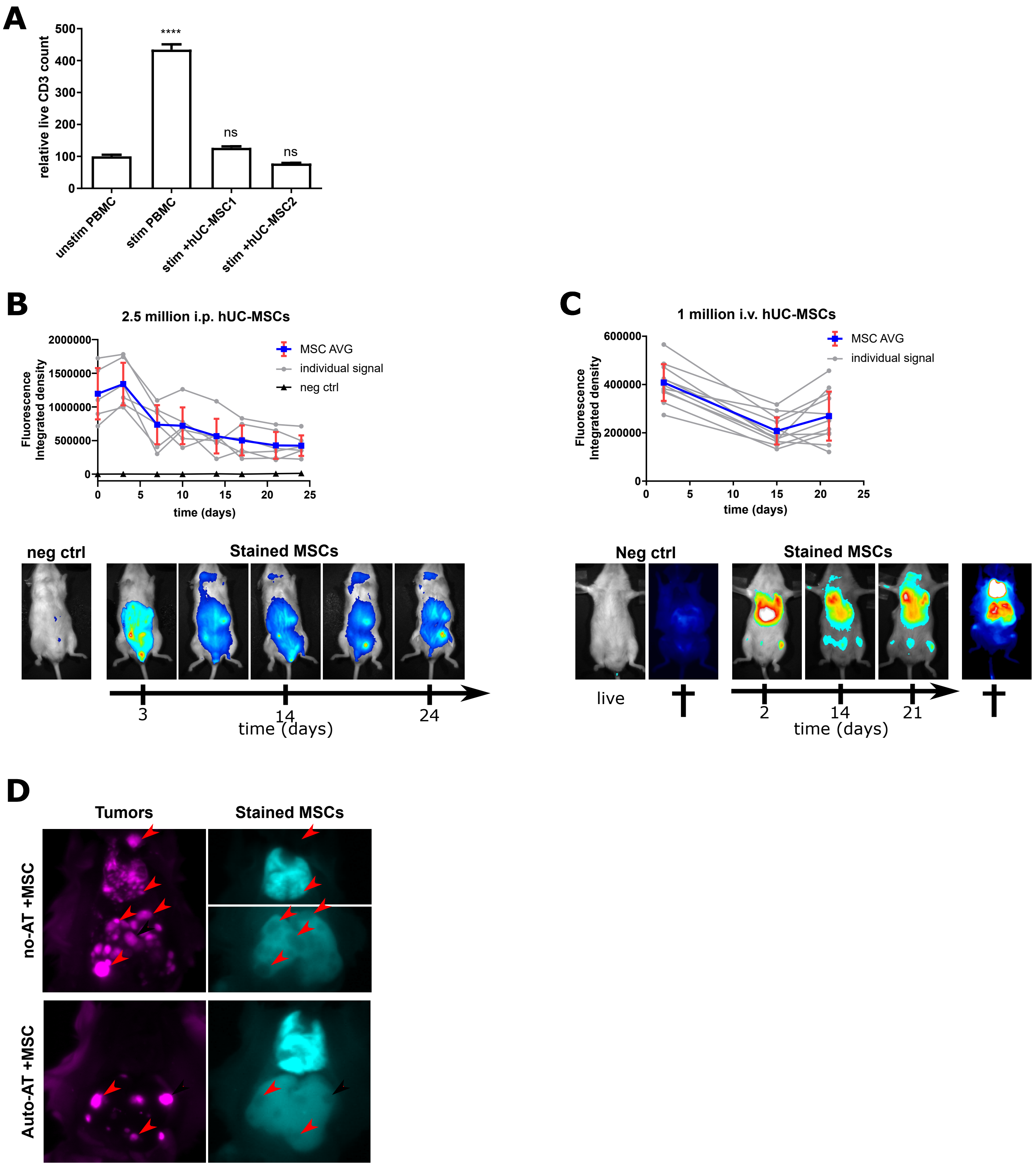


**Fig S3. hUC-MSC validation.** (A) *In vitro* hUC-MSC immunomodulatory potential assessment by T cell proliferation inhibition. Statistical analysis relative to unstim PBMC condition. (B) *In vivo* persistence of $2.5x10^6$ NIR fluorescent-stained hUC-MSCs after i.p. injection. Signal quantification (top) and sample images (bottom). Mean±SD. (C) *In vivo* persistence of $1x10^6$ NIR fluorescent-stained hUC-MSCs after i.v. injection. Signal quantification (top) and sample images (bottom). Mean±SD. (D) Tumor/hUC-MSC colocalization in metastatic-like tumor model. Tumor (left) and hUC-MSC (right) fluorescent signal with red arrowheads highlighting tumor mass positions in the absence (top) or presence (bottom) of autologous AT. Exposition for no-AT +MSC hUC-MSC signal in the lung and liver was adjusted independently for clarity.

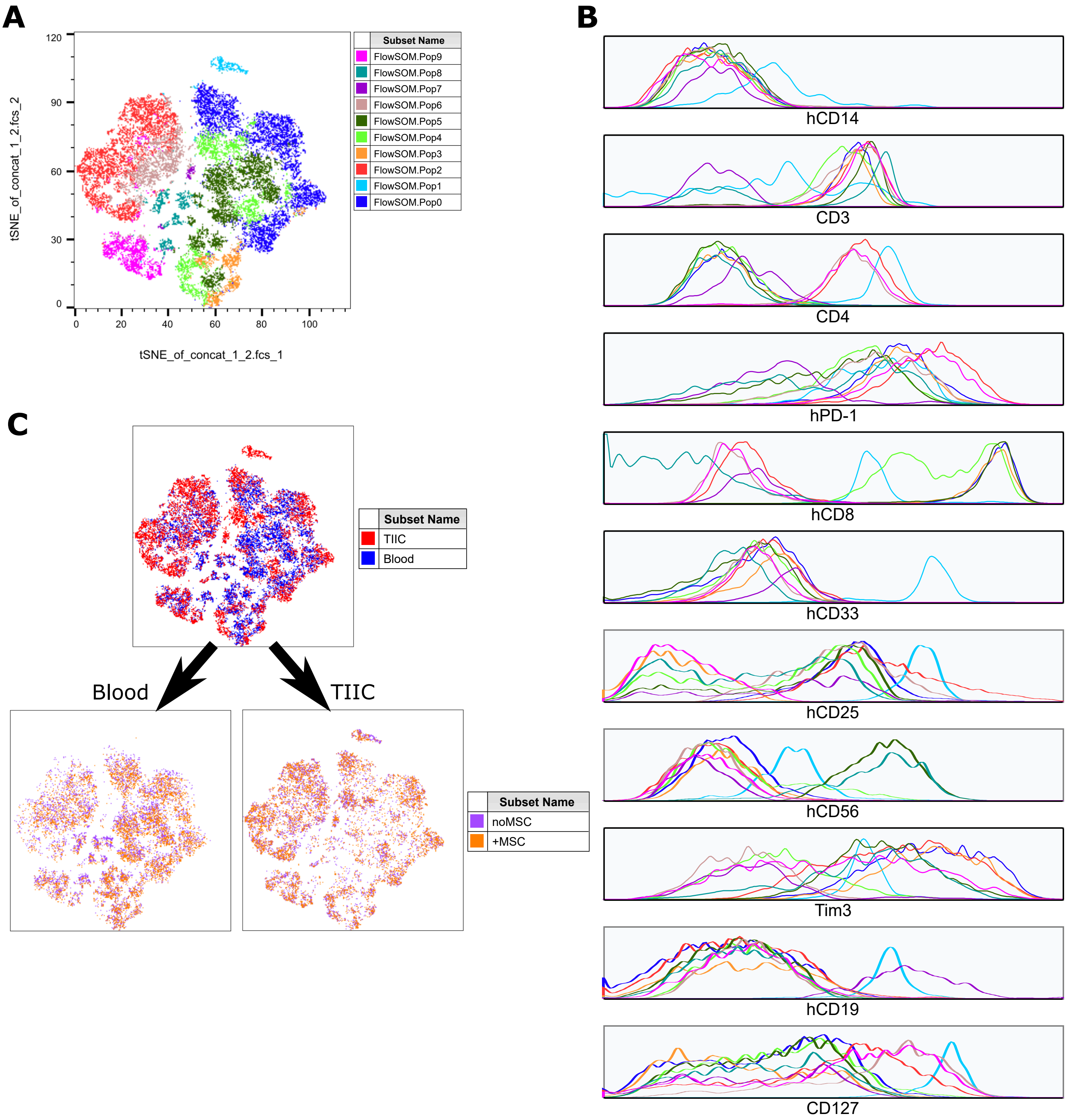


**Fig S4. Complementary gene expression profile for Auto-AT human lymphocyte clustering.** (A) Reference tSNE dimensional reduction plot of Auto-AT hCD45+ cells and (B) detailed gene expression profile for all population clusters generated using the FLOWSOM plugin in FlowJo (bottom). Each line color corresponds to its equivalent population in the tSNE plot. (C) Differential clustering between blood and TIICs (top) and differential clustering of TIICs with or without hUC-MSCs in blood (bottom left) or TIICs (bottom right). Signal overlap indicates no difference inpopulation distribution.

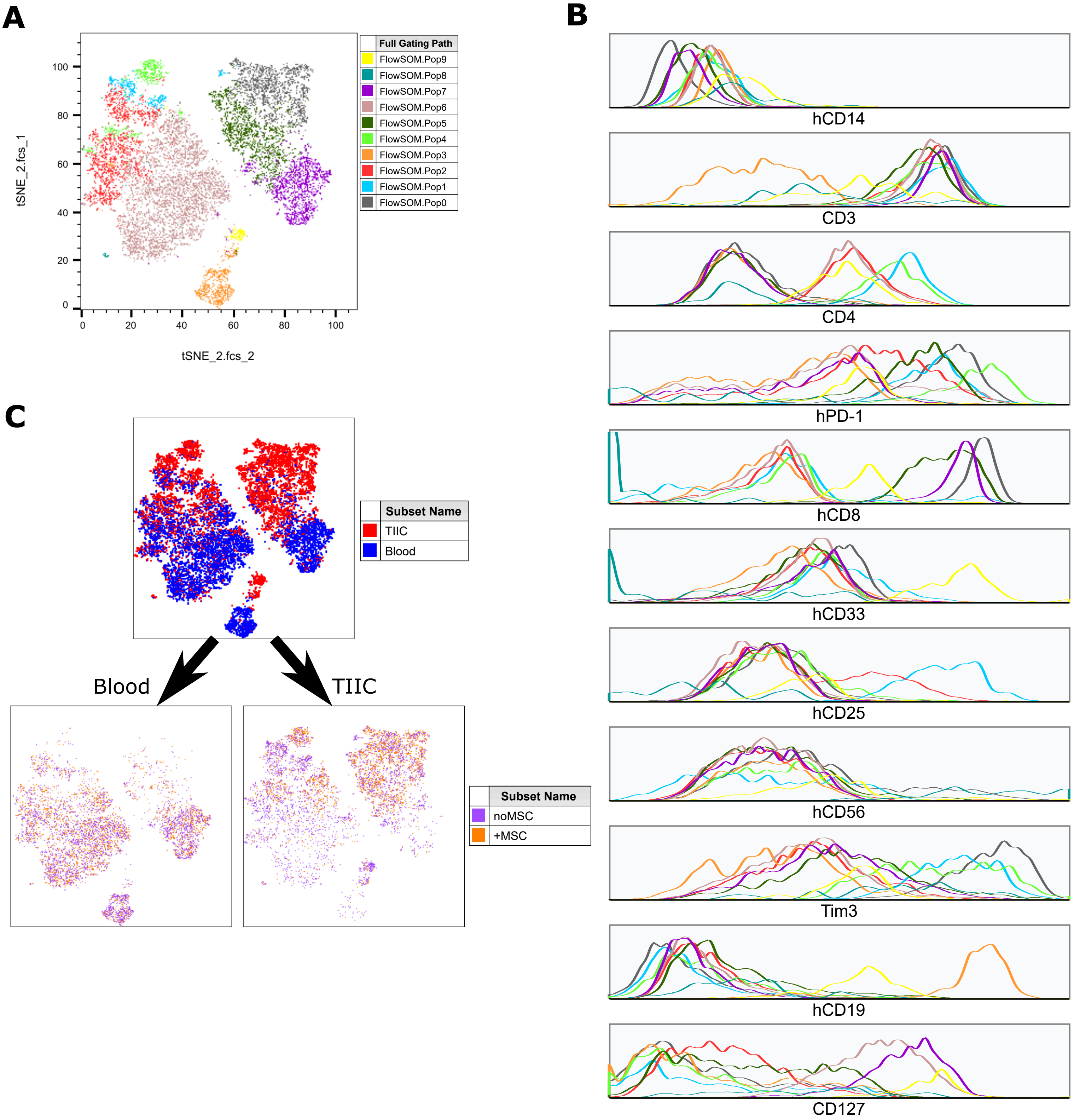


**Fig S5. Complementary gene expression profile for BLT human lymphocyte clustering.** (A) Reference tSNE dimensional reduction plot of BLT hCD45+ cells (top) and detailed gene expression profile for all population clusters generated using the FLOWSOM plugin in FlowJo (bottom). Each line color corresponds to its equivalent population in the tSNE plot. (C) Differential clustering between blood (blue) and TIICs (red) (top) and differential clustering of TIICs without (purple) or with hUC-MSCs (orange) in blood (bottom left) or TIICs (bottom right). Signal overlap indicates no difference in population distribution.
